# Regulatory mechanisms are revealed by the distribution of transcription initiation times in single microbial cells

**DOI:** 10.1101/223552

**Authors:** Sandeep Choubey, Jane Kondev, Alvaro Sanchez

**Affiliations:** Department of Physics, Brandeis University, Waltham, Massachusetts 02453; Rowland Institute at Harvard, Harvard University, Cambridge, Massachusetts 02142; Department of Ecology and Evolutionary Biology, Microbial Sciences Institute, Yale University, New Haven CT 06520

## Abstract

Transcription is the dominant point of control of gene expression. Biochemical studies have revealed key molecular components of transcription and their interactions, but the dynamics of transcription initiation in cells is still poorly understood. This state of affairs is being remedied with experiments that observe transcriptional dynamics in single cells using fluorescent reporters. Quantitative information about transcription initiation dynamics can also be extracted from experiments that use electron micrographs of RNA polymerases caught in the act of transcribing a gene (Miller spreads). Inspired by these data we analyze a general stochastic model of transcription initiation and elongation, and compute the distribution of transcription initiation times. We show that different mechanisms of initiation leave distinct signatures in the distribution of initiation times that can be compared to experiments. We analyze published micrographs of RNA polymerases transcribing ribosomal RNA genes in *E.coli* and compare the observed distributions of inter-polymerase distances with the predictions from previously hypothesized mechanisms for the regulation of these genes. Our analysis demonstrates the potential of measuring the distribution of time intervals between initiation events as a probe for dissecting mechanisms of transcription initiation in live cells.

## Introduction

One of the key findings of the genomic era is the unexpectedly high similarity between the genomes of different organisms(1). As the number of genomes being sequenced is increasing, it is becoming clear that the biggest difference among organisms is not to be found in their protein-coding sequences, but in the ways in which their genes are regulated(2–5). This is putting the spotlight on the parts of the genome that are responsible for gene regulation, and prompting the question of how changes in these sequences alter the way in which cells respond to intra and extracellular signals(6).

A large amount of genetic regulation occurs at the level of transcription, where cells control the amount of messenger RNA of each gene they express(7). Regulation of transcription is commonly achieved by the integration of multiple intracellular signals at the regions of DNA upstream from and in proximity to the gene’s coding region. This “promoter region” consists of a collection of transcription factor binding sites and, in eukaryotes, nucleosome positioning sites. Together, these binding sites dictate the binding and unbinding of specific transcription factors, co-factors and chromatin remodeling factors, which in turn either promote or inhibit the assembly of the transcriptional machinery at the gene.

The collection of transcription factor binding sites (that includes enhancer regions in eukaryotes), their position, and affinity for transcription factor proteins, constitutes the promoter architecture. Considerable effort has been directed to elucidating how promoter architecture determines measurable quantities like the average transcriptional response(8–11) of cells to a given stimulus, as well as the population-wide fluctuations of that response (12–15). While we have witnessed considerable progress on this front in recent years, many mysteries remain even in simple organisms such as bacteria. In particular, how promoter architecture affects the dynamics of transcription initiation in single cells remains poorly understood. To answer this question, experiments are being done where the number of RNA molecules from a gene of interest is measured at a single cell level in a population of isogenic cells (12, 16–18). The measured distribution of mRNA numbers in the cell population can then be used to test the predictions of different models of transcription initiation in the hope that some of these are supported by the data(19–30). This approach has led to the discovery of bursty mechanisms of transcription initiation (12, 16). However, this method of inferring the kinetics of transcription is limited by the fact that the mRNA copy number reflects additional processes downstream of transcription, such as the non-linear degradation of mRNA and proteins (31), maturation time of fluorescent reporters (32), mRNA transport (33), mRNA splicing (34) and small RNA regulation (35). The stochastic nature of these processes may introduce fluctuations in the number of mRNAs that masks the contribution of the transcriptional dynamics (36, 37).

In contrast, experiments that catch RNAP molecules in the process of transcribing a gene provide a more direct readout of transcription initiation dynamics. Techniques developed by O. Miller and his group in a series of landmark papers over several decades rely on imaging actively transcribed genes in recently lysed cells by electron microscopy (38–43). In these images, the positions of transcribing polymerases along a gene can be determined (Fig. 1A). Inter-polymerase distance distributions can be extracted from these positions and, with a few reasonable assumptions (36), these provide information about the distribution of times between successive initiation events (Fig. 1C). The information contained in these distributions of inter-polymerase distances is akin to that obtained in live cells by fluorescently labeling nascent RNAs to observe transcription initiation events in real time at the single cell level (19, 20, 44–48), as shown in Fig. 1B.

**Fig. 1.**
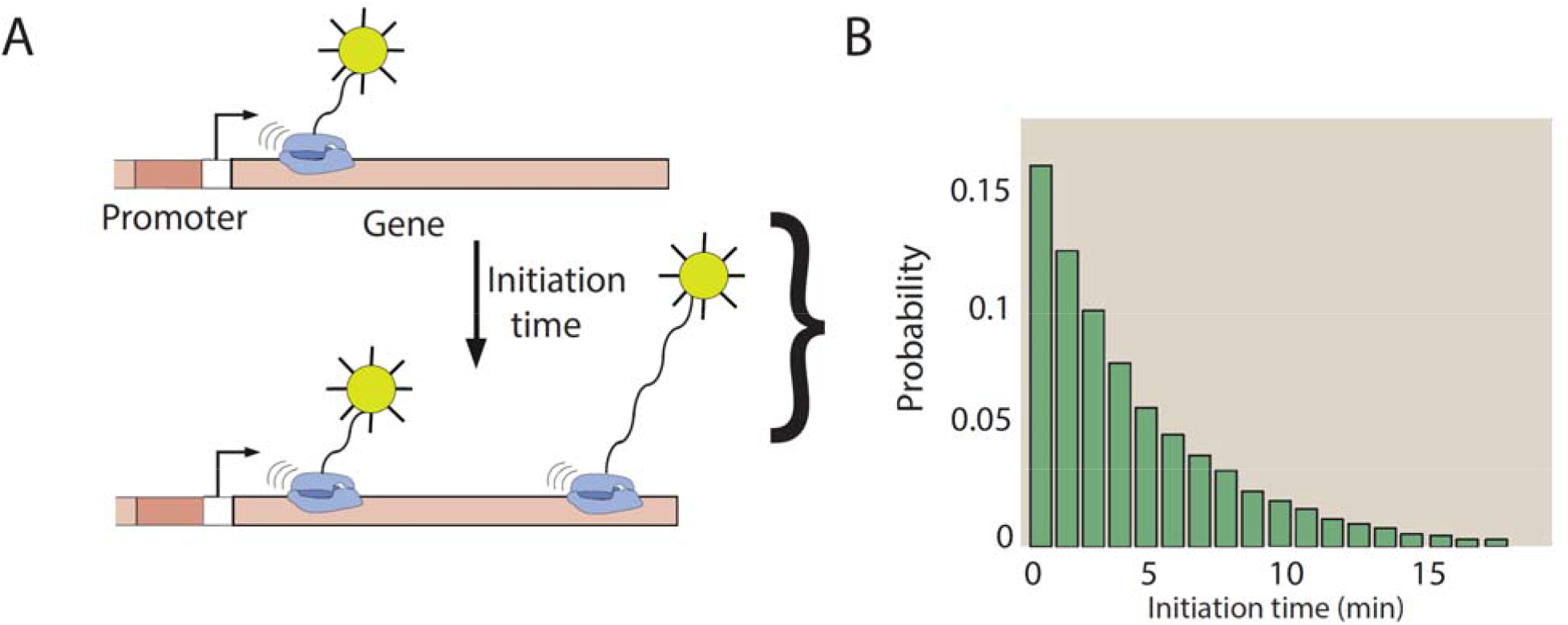
Positions of transcribing RNA polymerases carry the signature of transcription initiation dynamics. Schematic of the key idea of this paper. (A) Times between successive transcription initiation events can be extracted at the single cell level using fluorescent reporters for nascent RNA molecules (19, 20, 44–48), or from electron microscopy (EM) images of RNA polymerases caught in the process of transcribing a gene (38–43). Native elongating transcript sequencing (NET-seq) (80) can also obtain the same quantitative information as EM images. (B) The distribution of times between individual transcription initiation events can be extracted from experiments and compared to theoretical predictions based on stochastic models of transcription initiation.

Here we calculate the distribution of times between successive initiation events to quantitatively test mechanistic models of transcription initiation in cells. To accomplish this, we introduce a stochastic model of transcription that incorporates both initiation and elongation kinetics. Using the derived analytical results in conjunction with simulations, we show that the kinetics of initiation leave a signature in the distribution of transcription initiation times that can be used to discern between different models of transcription initiation. To showcase the power of this approach, we have re-analyzed a set of micrographs of *E.coli* genomic DNA, that, to our knowledge, provided the first reported evidence of transcriptional bursting, a phenomenon that was later found to be widespread across all organisms (49). We show that by filtering the information contained on these micrographs through our theoretical framework, we can test various proposed models of regulation of ribosomal promoters in *E. coli* in response to an increase in rrn operon copy number. We find that some of the previously proposed models produce distributions of inter-polymerase distances that are inconsistent with the Miller spread data, whereas others are in excellent agreement.

## Distribution of initiation times for an arbitrary mechanism of transcription initiation can be computed from a master equation

Transcription initiation is typically regulated by transcription factors and co-factors that bind to the regulatory DNA sequences and either inhibit or aid the binding of RNAP molecules to the promoter. To connect mechanisms of transcription initiation with measured times between successive initiation events, we consider a stochastic model of transcription with a general initiation mechanism, where the promoter can be in an arbitrary number of states defined by different constellations of bound transcription factors and co-factors. Using a chemical master equation approach (22, 50, 51), we show that the distribution of the distribution of times between two initiation events and its moments can be computed analytically for any mechanism of transcription initiation. These equations allow us to discriminate between different mechanisms of initiation by comparing the predicted distributions to experimental distributions of transcription initiation times.

In order to compute the distribution of times between successive initiation events, we assume that the promoter can be in *N* different discrete biochemical states and that transitions between different states occur as different transcription factors bind and fall off their respective binding sites. The rate of transition from the *m*-th to the *n*-th promoter state is *k_m,n_*, and the rate at which an RNAP molecule initiates transcription from the *m*-th promoter state is *k_m,esc_*. The assumption that the transitions between these states are random Poisson processes characterized by rate constants leads to a chemical master equation that describes the time evolution of *P_m_(t)*, the probability that the promoter is in the *m*-th state (*m = 1, 2,…, N*) at time t:

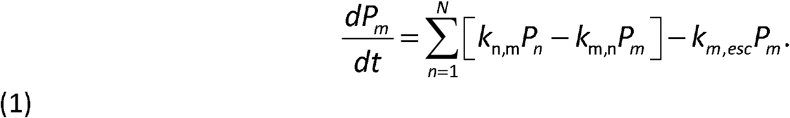

Solving this chemical master equation for all the states *m* from which the promoter initiates transcription with rate *k_m,esc_*, leads to a general formula for the probability distribution *q*_1_(*t*) of the time intervals between successive transcription initiation events: (details of the calculation can be found in the SI)

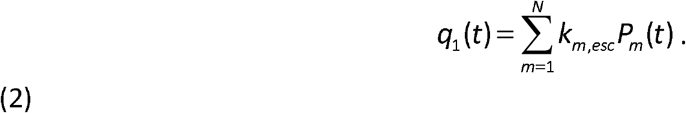

Assuming a uniform elongation rate (37, 52) along the gene *ν*, the distribution *q*_1_(*t*), of time intervals between successive initiation events directly translates into a distribution of distances *q*_1_(*x*) between RNAPs along the gene. In other words, the inter-polymerase distance distribution along a gene is given by,

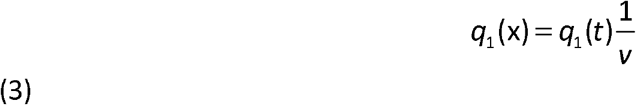

In the ensuing investigation of regulation of ribosomal genes, Equation 3 forms the basis of our analysis of positions of RNAP molecules along a gene at a given moment in time, which provides a quantitative test for different models of transcription initiation. Although transcription elongation of ribosomal genes is typically more complicated and involves pausing and backtracking of polymerases along the gene (53), here we assume that transcriptional pausing happens on time scales that are negligible compared to the times between transcription initiation events(54). For a detailed discussion of this model assumption, see the SI.

## The distribution of transcription initiation times can be used to discern between different models of initiation

To illustrate how the distribution of times between successive initiation events can be used to extract mechanistic insights into the process of transcription initiation, we consider three different models of initiation as case studies (see Fig. 2).

**Fig. 1.**
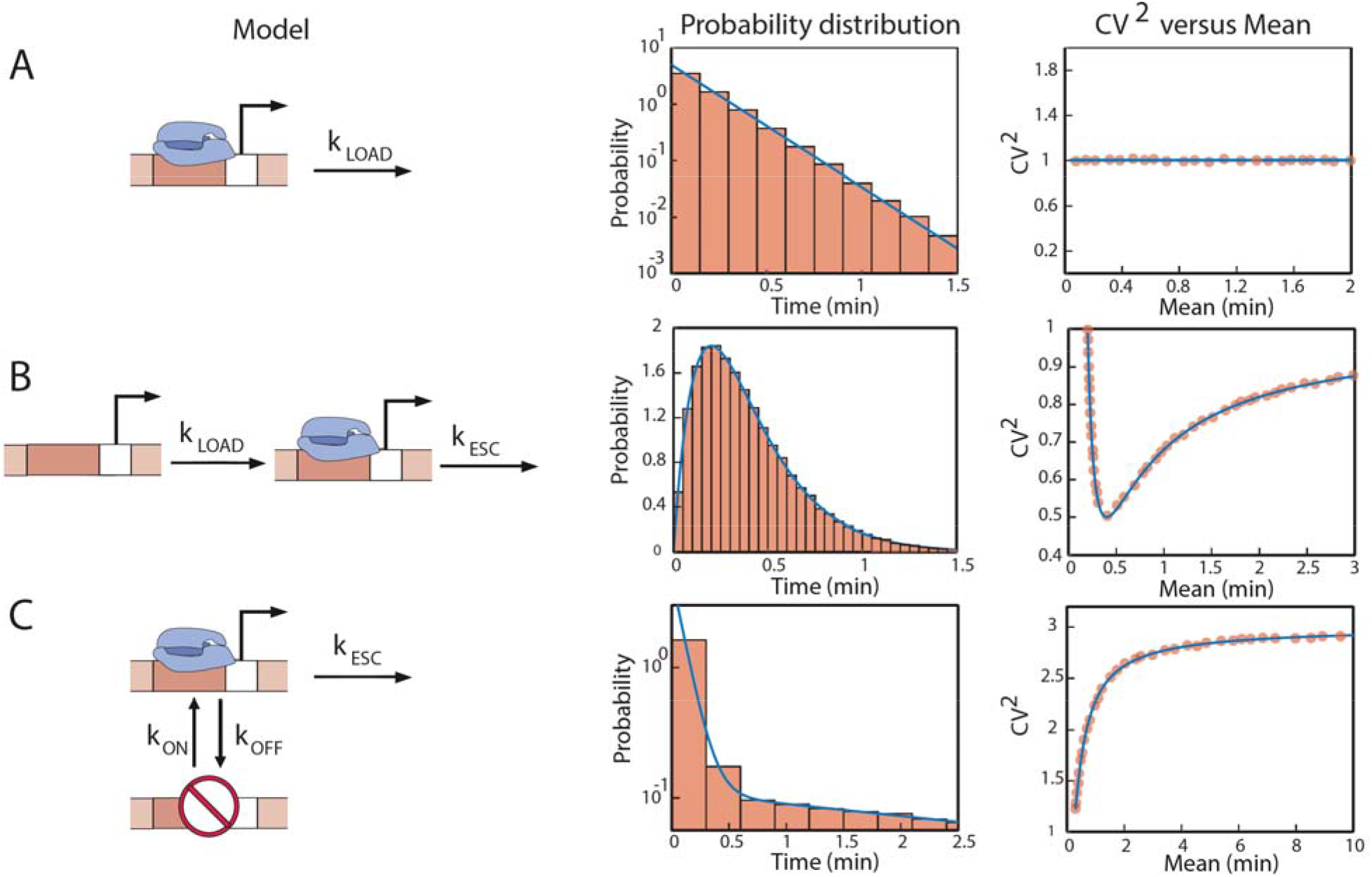
Different models of transcriptional regulation leads to distinct signatures in the initiation times: (A) One-step model of transcription initiation. Initiation happens at a constant rate *k_LOAD_*. The times between successive initiation events are exponentially distributed. The square of the coefficient of variation is plotted as a function of the mean, where we change the mean by changing the rate of initiation, *k_LOAD_*. We confirm the analytical results using Gillespie simulations(61). The histogram and closed circles represent simulation results. (B) Two-step model of transcription initiation. Initiation happens in two sequential steps: the rate of RNAP loading on to the promoter occurs with rate *k_LOAD_*, followed by RNA polymerase escaping the promoter leading to initiation event at a rate, *k_ESC_*. The distribution of times between successive initiation events and the square of the coefficient of variation of the distribution, as a function of the mean are shown. To change the mean, we change the rate of loading of RNAP polymerase molecules on the promoter, *k_LOAD_*. As before, simulation results are compared to the analytical results. (C) ON-OFF model: The promoter switches between two states: an active and an inactive one. The rate of switching from the active state to the inactive state is *k_OFF_*, and from the inactive to the active state is *k_ON_*. From the active state transcription initiation proceeds with a probability per unit time, *k_ESC_*. The distribution of times between initiation events, and the square of the coefficient of variation as a function of the mean are shown. Results from Gillespie simulations(61) are shown for comparison. To change the mean, we tune the rate *k_ON_* of switching from the inactive to the active state. To illustrate the distinctive impact of the different initiation models on the distribution and moments of the times between successive initiation events, we use the following parameters: *k_OFF_*=5/min, *k_ON_*=0.435/min, *k_LOAD_*=0.14/min and *k_ESC_*=0.14/min, which are characteristic of yeast promoters(36).

### Poisson (single rate limiting step) model

The Poisson model is the null model of initiation, which is usually associated with constitutive promoters (12). In this model initiation happens with a constant probability of *k_LOAD_* per unit time, as shown in Fig. 2A. In bacteria, this step could, at the molecular scale, represent the rate of loading of RNAP molecules to the promoter DNA, while for eukaryotes this step could correspond to the formation of the pre-initiation complex. As obtained from Eqn. 2, the one-state model is characterized by exponentially distributed times between successive initiation events. One of the key properties of an exponential distribution is that its mean and standard deviation are equal. Therefore, the CV^2^(defined as the ratio of the variance to the square of the mean) is always equal to one, independent of the rate *k_LOAD_*, as shown in Fig. 2A.

### Two limiting steps model

Next, we consider a model in which initiation happens in two sequential rate limiting steps. This is the situation when two steps in the sequence of events leading to initiation are of comparable duration. For example, in bacteria the first step could correspond to an RNAP molecule binding to the promoter with a rate *k_LOAD_*. In Eukaryotes this step could represent the loading of the transcriptional machinery at the promoter (23, 47). In the second step, the promoter bound RNAP molecule escapes the promoter at a rate *k_ESC_*, and starts transcribing the gene. For several promoters in yeast(36) and *E.coli* (55), it has been reported that initiation proceeds through two-sequential steps.

For this case, using Eqn.2, we find that the waiting time distribution between successive initiation events is gamma distributed. This result agrees with previous theoretical studies(25) and it leads to the following relationships between the kinetic rates of the mechanism (*k_LOAD_* and *k_ESC_*) and the mean and the coefficient of variation of the waiting time distribution:

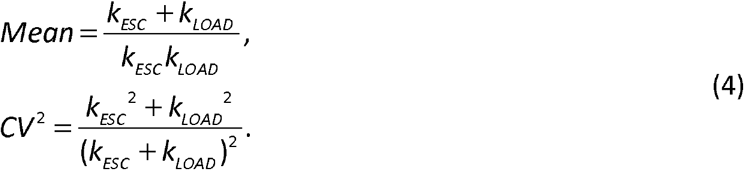

As shown in Fig. 2B, when we tune either one of the two rates of the model while keeping the other one constant, the coefficient of variation initially decreases as a function of the mean, develops a minimum when the two rates become equal, and then asymptotically goes to one. In the limit of one rate being much slower than the other one, the waiting time distribution becomes exponential, leading to a coefficient of variation of one.

### ON-OFF promoter

The third scenario we consider is the ON-OFF model of initiation. This model of initiation has been established as the canonical model of transcriptional regulation for both bacteria (19) and eukaryotes (17, 56–60). In this model, the promoter switches between two states: an active state, from which transcription initiation can occur, and an inactive state from which initiation does not occur. The two states might correspond to a free promoter and one bound by a repressor protein, or a promoter occluded by nucleosomes. The rate of switching from the active to the inactive state is *k_OFF_* and from inactive to the active state is *k_ON_*. The rate of initiation from the active state is *k_ESC_*.

In this case we find that the waiting time distribution between successive initiation events is given by a sum of two exponentials, as shown in Fig. 2C. Thus, it can be distinguished from a single exponential expected from the one-state promoter, on the condition that the decay constants of the two exponentials are well separated in magnitude. The mean and the coefficient of variation as functions of the different biochemical rates are given by,

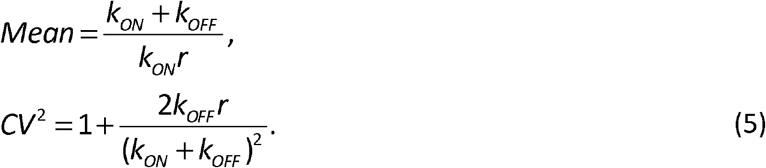

When we tune the rate *k_ON_*, the CV^2^ increases as a function of the mean and eventually saturates, as shown in Fig. 2C.

We compare our analytical results for the three models described above against Gillespie simulations(61). This allows us to numerically generate multiple time-traces of initiation events. From these different time traces, we obtain the distribution of initiation times as well as the corresponding values of the mean and variance. The histograms for the times between initiation events for all the three models are shown in Fig. 2A-C. We also show the coefficient of variation as a function of the mean for these models, as we tune the relevant rates in Fig. 2A-C. These results imply that we can discern these three different models of initiation based on the predictions they make for the waiting time distribution of consecutive initiation events as a function of the different experimentally tunable parameters.

## The dynamics of transcription initiation of ribosomal genes in *E.coli* can be extracted from images of transcribing polymerases in fixed cells

To demonstrate how the distribution of inter-polymerase distances along a gene can be used to extract dynamical information about the process of transcription initiation *in vivo*, we have reanalyzed images of elongating RNAP molecules on ribosomal RNA (rRNA) genes in *E. coli*, which were obtained using the Miller spread technique by Voulgaris et al. (38). In Fig. 3C, we show the inter-polymerase distance distribution for the seven ribosomal genes in wild type *E. coli* cells (strain pBR322 (38)). A remarkable feature of this distribution is the presence of a peak in the inter-polymerase distance distribution at small distances. This is inconsistent with a “Poissonian” initiation mechanism (19). Indeed, the presence of a maximum in the probability at intermediate distances suggests the existence of a two limiting steps model of initiation (Fig. 3B), where, for example, the polymerase first binds to the promoter, and then escapes the promoter leading to elongation, where the two steps occur with comparable rates. Recent *in vivo* studies in yeast have shown that initiation can proceed in multiple-sequential steps, where the rates involving these steps have comparable magnitude (36). In addition, when analyzing this data, we consider the time it takes for the polymerase to clear the promoter by elongating through it. To test the hypothesis of two sequential steps leading to initiation, we fit Eqn.3 to the experimental distribution obtained from the images, while assuming as a parameter an elongation speed of 78 bp/second (as measured elsewhere (38)). We find that the two limiting steps model is in good agreement with the data. Furthermore, the fit provides estimates for the rates of promoter escape, rate of RNAP loading on to the promoter, and time to clear the promoter (*k_LOAD_* ≈ 3/second, *k_ESC_* ≈ 3/second and *τ_CLEAR_* ≈ 0.3 seconds), all of which are in good agreement with previous measurements (62) (See the Materials and Methods section).

**Fig. 3:**
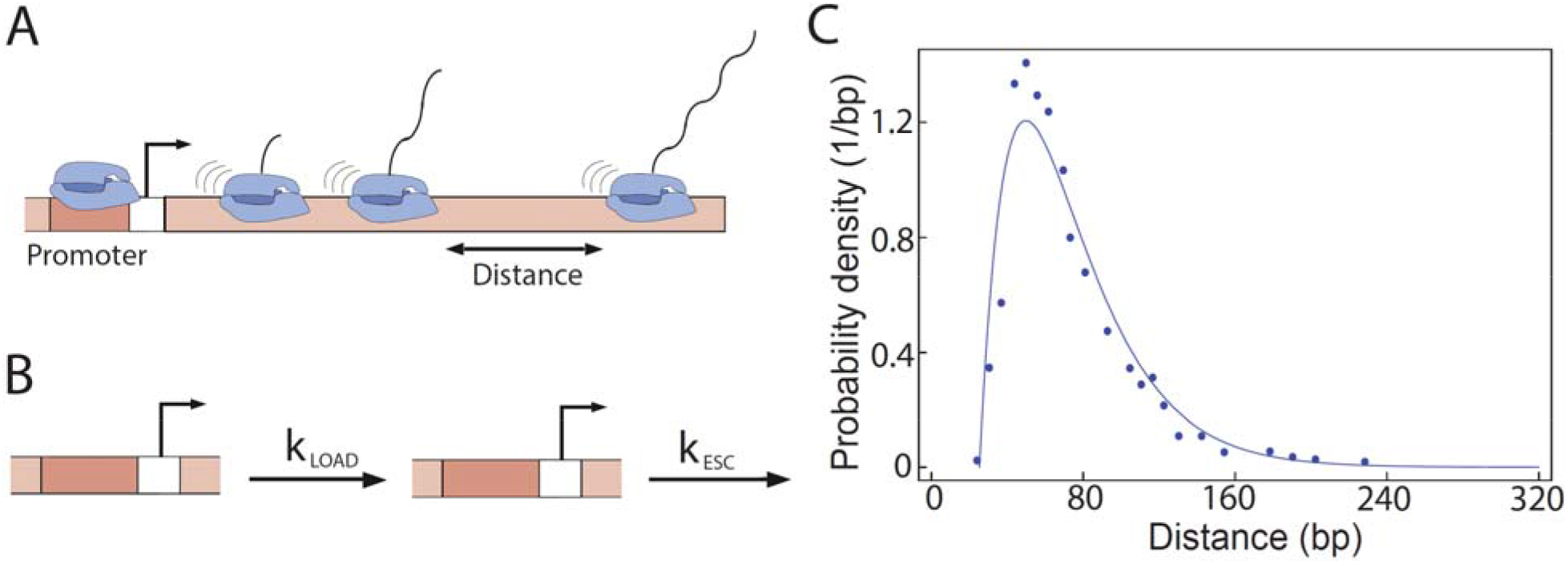
Initiation of transcription of ribosomal genes in *E.coli*: (A) Positions of RNA polymerase molecules transcribing a gene at a given instant in time can be obtained from electron microscopy images or native elongating transcript sequencing (80). (B) Two-step model of transcription initiation, as shown in Fig. 2B. (C) Fit (line) of the two-step model to the inter-polymerase distance distribution data (points) obtained by Voulgaris et al. (38) for ribosomal genes in *E.coli*. The different biochemical rates we extract are *k_ESC_* (rate of promoter escape)≈3/second, *k_LOAD_* (rate of RNAP loading on to the promoter)≈3/second and *τ_clear_* (time for a RNAP to clear the promoter) ≈ 0.3 seconds, taking the elongation speed v= 78 bps/sec, as reported in experiments(38).

## Transcriptional bursting accompanies the down-regulation of the expression of rrn operons in the presence of additional copies of the rrn genes

In a second set of experiments, electron micrographs images were used to shed light onto a previously reported effect (63), namely that the transcriptional activity of individual rrn genes is inversely proportional to the copy number of these genes, in such a way that the net transcriptional output of rrn genes in the cell is kept constant. Surprisingly, the electron micrographs showed that when the copy number of ribosomal genes was altered by placing extra rrn genes on plasmids, a very different pattern of polymerase occupancy of the rrn genes emerged. Beside the mean number of RNAP molecules along each gene decreasing with increased gene copy number, it was also observed that RNAP molecules are now grouped in bunches along the gene, which is indicative of transcriptional bursting. Bremer et al.(64) proposed that since a significant fraction of RNAPs is engaged in transcribing the ribosomal genes, then changing the gene copy number will significantly alter the concentration of free polymerases available for transcription initiation. This will consequently decrease the rate of transcription initiation, which is assumed to be proportional to the RNA polymerase concentration. This “free RNA polymerase hypothesis” (64) along with the two-step model of transcription initiation (Fig. 1B) predicts that the mean number of RNAPs per gene will decrease in response to an increase in the number of genes, as observed experimentally, but it cannot reproduce the observed bunching of polymerases, as was shown in a previous theoretical study (25). Therefore, in order to account for the bunching of RNAPs seen in experiments by Voulgaris et al. (38), it is necessary to consider models of initiation with promoter states that are off-pathway to elongation. Indeed, several such models have been proposed to explain the down-regulation of individual rrn genes in response of an increase in the gene copy number(38). These models can be broadly classified into three different classes. Two of these three classes of models are extensions of the two-step model to include off-pathway promoter states that may lead to bursting of transcription initiation which would then lead to gaps between bunches of RNAP molecules along the gene. Below we test the predictions of these three classes of models for the distribution of initiation times against the experimental data.

The first class of models, considers the formation of long-lived non-productive initiation complexes at the promoter (65–67). For example, non-productive complexes that cannot exit the abortive initiation state into productive elongation have been observed *in vitro* (25). The formation of such dead-end complexes can block the promoter for long periods of time. This promoter blockade leads to transcriptional bursting which could in turn produce the gaps between bunches of RNAPs transcribing the gene, shown in Fig. 3A.

The second class of models, assume cooperative recruitment of RNAP molecules to the promoter by a RNAP molecule already present on the promoter (68–70). For example, when a RNAP molecule initiates transcription it can leave the promoter DNA in a supercoiled state as illustrated in Fig. 3B. In the supercoiled state, the energy barrier for melting a strand of DNA to make a transcriptional bubble is lowered leading to an increased rate of RNAP loading on to the promoter (25). If the rate at which promoter DNA relaxes from the supercoiled state is not much larger than the RNAP loading rate, then several polymerases can initiate in a burst of activity leading to the formation of a bunch of RNAPs along the gene. When the promoter DNA relaxes from the supercoiled state, the rate of loading of polymerase molecules, which leads to the creation of a gap between successive RNAP bunches. The kinetic steps of this model are shown in Fig. 3B. It has been proposed by several authors that negative supercoils introduced by RNAP initiating transcription may induce such cooperative recruitment (25). In fact Voulgaris et al. (38) in their paper speculated that a possible reason for the observed gaps in the distribution of RNAPs along the gene could be supercoiling-mediated recruitment of RNAPs.

A common feature of both classes of models is that they incorporate the “free RNAP” hypothesis, in that an increase in the number of rrn genes in the cells leads to a reduction in the rate of loading of RNAP molecules to the promoter. In contrast, a third alternative class of regulatory models for the transcription of ribosomal genes have been proposed, whereby a secondary molecular messenger whose abundance in the cell is regulated in response to the changing number of ribosomal genes either inhibits or activates the transcription of these genes (Fig. 3C) (71). One such model is based on the alarmone nucleotide molecule ppGpp, which is capable of inactivating the promoter-RNAP complex upon binding to the polymerase (71, 72). The inactivated polymerase effectively blocks further transcription and leads to the appearance of gaps between bunches of transcribing polymerases, as observed in the micrographs of Voulgaris et al. (38). The key assumption of this model that distinguishes it from the first two is that an increase in the number of rrn genes does not significantly change the number of free RNAPs in the cell (71, 72).

Using our mathematical framework, we put to test these different classes of models, with the goal of gaining insight into the unresolved question of how ribosomal genes are regulated when the number of gene copies is increased. To do so, we compute the distribution of distances between transcribing polymerases based on these models of initiation using Eqn. 3, and then compare the results directly with the distribution obtained from electron-microscopy images. In order to analyze the inter-polymerase distance data, Voulgaris et al. (38) treated the polymerase distances within a bunch and between bunches separately. Hence to compare our theory with the experiments, we compute the mean and the variance of the distances between polymerases within a bunch and between bunches for all three models. It is important to note that despite the ribosomal genes being highly transcribed their inter-polymerase distance distributions are not significantly affected by the elongation dynamics (for a detailed discussion of this point see the SI).

We find that these different classes of models, as shown in Fig. 4, make starkly different predictions for the inter-polymerase distributions along the gene. When the computed distributions are compared to those measured in experiments, we find that the third class of models, where a secondary molecular messenger inhibits the transcription of the gene is favored over the two models that are based on the “free RNAP” hypothesis. Although the two “free RNAP” classes of models may produce bursts of transcription initiation, the theoretical predictions these models make for the intra-bunch mean and variance are inconsistent with the data obtained by Voulgaris et al. (38), (Fig. 4A,B). A common feature of the “free RNAP” models is that they predict an increase in the mean and the variance in the distribution of RNA polymerase distances within a “bunch”when the number of ribosomal genes is increased. This stems from the assumption that changing the gene copy number leads to a lower concentration of free RNAP, and thus to a reduction in the rate of RNAP loading on to the promoter (red arrow in Fig. 4A,B). However, when the experimental distribution of initiation times we found that increasing the number of genes from seven to ten has no significant effect on the distribution of distances between RNAPs within a bunch. This is inconsistent with the theoretical prediction from these two mechanisms.

**Fig. 4:**
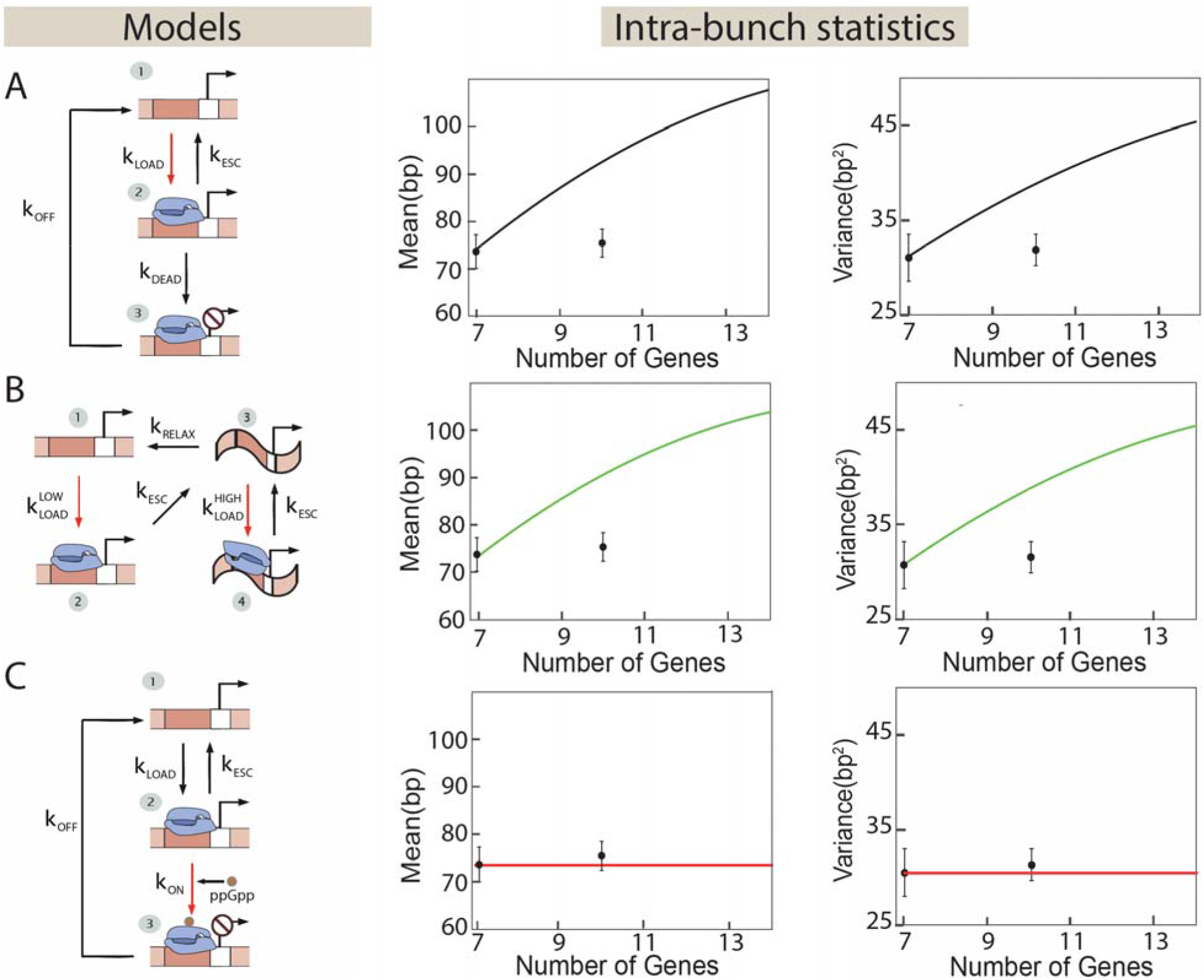
Different models of transcriptional regulation of ribosomal genes can be tested by tuning the gene copy number. (A) and (B) Models of transcription initiation that rely solely on the interaction of RNA polymerases with promoter DNA. (A) This class of model considers the formation of long-lived non-productive initiation complexes at the promoter by RNAP molecules (25, 67). After binding the promoter at a rate *k_LOAD_*, each RNAP can initiate transcription at a rate *k_ESC_* or make a dead-end complex at the promoter at a rate *k_DEAD_*. These dead-end complexes are unproductive and are removed at a rate *k_OFF_*. The change in gene copy number affects the binding rate of RNAP molecules to the promoter due to a change in the free RNAP concentration, as indicated by the red arrow. Theory predicts that the mean and variance of distances between RNAPs within a bunch increase with the gene number, contrary to experiments on ribosomal genes in *E.coli*. (B) Cooperative recruitment of RNAP by DNA supercoiling. RNAP molecules are loaded on to the promoter at a rate *k_LOAD_^LOW^*. After RNAP initiates transcription, at a rate *k_ESC_*, it leaves the promoter DNA in a supercoiled state and subsequent loading of RNAP polymerases at the promoter at a faster rate *k_LOAD_^HIGH^*. The rate of relaxation of the supercoiled state is *k_RELAX_*. The change in gene copy number affects both the polymerase loading rates (red arrows) due to the change in free RNAP concentration. The model predicts that the mean and variance of the intra-bunch RNAP distances increase with the gene copy number, contrary to measurements in *E.coli*. (C) As the number of genes increases the rate of ribosomal RNA production increases. This triggers the production of ‘control molecules’ (e.g. ppGpp) which then reduce the initiation rate by modulating the promoter-RNAP interactions. ppGpp regulates the initiation process by converting the active promoter-RNAP complexes into inactive ones. It is described by the same kinetic scheme as the dead-end complex model (A) with a critical difference, namely in this case it is the rate of ppGpp binding to the RNAP-promoter complex (red arrow) and not the rate of RNAP loading on to the promoter is tuned as the copy number of ribosomal genes is changed. Here we call the rate of inactivation of RNAP-DNA complex *k_ON_*. The mean and variance of distances between RNAPs within a bunch is predicted to remain constant as observed in experiments. In all the plots, the two data points shown are taken from (38).

In contrast, here we propose a third class of model based on the action of the alarmone nucleotide ppGpp. Recent experiments have shown that ppGpp molecules interact with active RNAP-promoter complexes and turn them into inactive ones(71). Formation of such inactive complexes can lead to bursty transcription initiation dynamics, caused by the resident polymerase blocking the promoter. Another set of experiments has shown that the number of ppGpp molecules increases with increasing ribosomal gene numbers (73). Consistent with these observations, we propose a kinetic model of transcriptional regulation of ribosomal genes where RNAP-promoter complexes are inactivated by ppGpp molecules that increases in concentration as the gene copy number is increased (shown in Fig. 4C). We computed the mean and variance of the inter-polymerase distance distributions within a bunch for this mechanism, and find that they are consistent with the experimental results. Since the key assumption of this model is that an increase in the number of genes does not significantly change the number of free RNAPs in the cell (71, 72), the rate of RNAP loading on to the promoter remains unchanged. This has the effect of keeping the distribution of distances between RNAPs within the bunch unchanged (Fig. 4C). Thus, the proposed mechanism generates the same bursting kinetics that has been found in electron micrographs for this promoter, suggesting that it may be a candidate explanation for the observed bursting pattern. We note that any mechanism based on stochastically and transiently preventing promoter escape by a bound polymerase would have the same outcome. While ppGpp is a plausible candidate as this mode of action has been documented before, whether it is acting through that mechanism here is still an open question.

## Discussion

The dynamics of transcription in live cells is poorly understood. Due to the difficulties in directly imaging the process of transcription (19, 46, 47, 49, 74), experimental methods for counting the products of transcription (such as RNA and protein molecules) in single cells have been developed over the past years. The protein and mRNA distributions carry the signature of the dynamics of transcription and hence can be exploited to decipher the underlying mechanisms of transcriptional regulation (12, 16, 46, 75). However both mRNA and protein counts are affected by noisy processes other than transcription such as mRNA processing, binomial partitioning, nonlinear degradation of mRNA molecules etc. (31, 33, 34, 76–79), which can potentially mask the signature of transcription on protein and mRNA distributions.

Recent experimental advancements make it possible to extract the positions of RNAP molecules that are engaged in the process of transcribing a gene at a given instant in time (38, 41–43, 80). Similar information can be extracted by observing transcription initiation events in real time using fluorescent reporters (19, 20, 44–48). These measurements are not affected by post-transcriptional processes and are therefore more direct readouts of transcription compared to mRNA and protein counting (81). In this paper, we have derived mathematical equations that allow us to interpret and analyze the inter-polymerase distance distribution, or equivalently, the waiting time distribution between successive initiation events across a population of isogenic cells. To demonstrate the potential utility of our analytical results, we fit the inter polymerase distance distribution for ribosomal genes in *E.coli* (acquired from electron micrographs) to a theoretical distribution computed for a two-step model of initiation. The model fits the data well, allowing us to extract the rates that characterize transcription initiation dynamics. We also reanalyze images of RNA polymerases transcribing ribosomal genes in wild type and mutant strains of *E.coli*. We show that previously proposed mechanisms, based on the effect of extra rrn gene copies have on the concentration of free RNAPs in the cell, are inconsistent with the observed inter-polymerase distance distributions. In contrast, we find that an alternative possibility(71), where the alarmone nucleotide ppGpp may interact with promoter bound RNAP and prevent promoter escape, produces bursting kinetics that are consistent with experimental observations. We believe that the approach and ideas presented here will be helpful to uncovering detailed, kinetic information about the process of transcription initiation in live cells.

## Methods section on

### Data analysis and parameter estimation

We evaluate the utility of our theoretical framework by employing it to gain mechanistic insights into the regulation of ribosomal genes in *E. coli*. To this end, we have re-analyzed a set of images of elongating RNAP molecules on ribosomal RNA (rRNA) genes in *E. coli*, which were obtained from electron micrographs of fixed cells using the Miller spread technique by Voulgaris et al. (38). To extract the digitized data from the plots (38), we use a software called, Digitizelt which is easily available online. The authors increased the number of rrn operons in *E. coli* cells by inserting an rrn operon on a multicopy plasmid. It was observed that the rate of rRNA expression per operon is reduced to maintain a constant number of rRNA in the cell. In fact, EM images showed that fewer RNAP molecules were engaged in transcribing the rrn genes, consistent with previous studies(63). Moreover, RNAP molecules formed bunches separated by gaps along the genes. While the authors ruled out transcription elongation or termination as origins of these bunches, they suggested that the bunches are caused by stochastic interruptions of initiation or promoter-proximal elongation events. The authors analyzed the EM images by defining a “transcriptional bunch” as a group of RNAPs separated by less than 240bp from each other (38). The distribution of distances greater than 240bp is referred to as inter-bunch distribution. Using our theory, we analyze the intra-bunch distributions.

To extract the model parameters for the dynamics of transcription initiation of ribosomal genes, we first consider the inter-polymerase distance distribution data for wild type *E. coli* cells. In Fig. 3C, we show the inter-polymerase distribution for the seven rrn promoters for the ribosomal genes in wild type *E. coli* cells (strain pBR322). We model the RNAP distance distribution within a bunch by taking the two-step model, shown in Fig. 3B. Using Eqn. 3 we find the probability distribution of inter-polymerase distances is given by

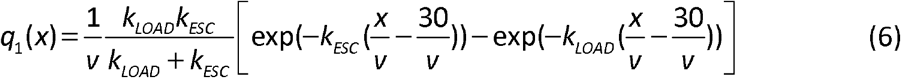

Here 30bps is roughly the size of a RNAP molecule (82). By comparing the data from the experiments for inter-polymerase distances within a bunch and the prediction from the model we extract *k_LOAD_*, *k_ESC_* and *τ_CLEAR_*. By fitting the model (see Fig. 3C), we extract the rates *k_ESC_* ≈ 3/second, *k_LOAD_* ≈ 3/second and *τ_clear_* ≈ 0. 3 seconds 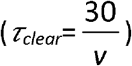, *v* is the rate of transcription elongation for every RNAP molecule, which we take to be 78 bps/sec (38).

Next, we seek to decipher the dynamics of transcriptional regulation of ribosomal genes by considered three models, as shown in Fig 4, as proposed by Voulgaris et al. (38). Each of these three models is defined by five parameters (see Fig. 4A,B and C). For the dead-end complex model, after the loading of polymerase molecules to the promoter at a rate *k_LOAD_*, each RNAP molecule either escapes the promoter at a rate *k_ESC_* and starts transcribing the gene, or forms a dead-end complex at the promoter at a rate *k_DEAD_*. These dead-end complexes are unproductive and are removed at a rate *k_OFF_*. For the Cooperative recruitment of RNAP by DNA supercoiling, RNAP molecules are loaded on to the promoter at the promoter at a rate *k_LOAD_^LOW^*. After RNAP initiates transcription, at a rate *k_ESC_*, it leaves the promoter DNA in a supercoiled state and subsequent loading of polymerases occurs at the promoter at a faster rate *k_LOAD_^HIGH^*. The rate of relaxation of the supercoiled state is *k_RELAX_*. For the third model, the production of ‘control molecules’ (e.g. ppGpp) reduce the initiation rate by regulating the initiation process by converting the active promoter-RNAP complexes into inactive ones. It is described by the same kinetic scheme as the dead-end complex model. However, the rate of inactivation of RNAP-DNA complex is given by *k_ON_*, every other rate remaining the same. To test these proposed models based on the experimentally observed transcriptional bunching data for rrn genes, we first extract the different parameters involving these models by fitting the inter-RNAP distance distributions these models produce with the data for wild-type *E. coli* cells with seven rrn operons. As demonstrated earlier, the intra-bunch distance distribution allows us to obtain three of the parameters, which are common to these models of initiation i.e. *k_LOAD_*(*k_LOAD_^HIGH^* for supercoiling mediated recruitment), *k_ESC_* and *τ_clear_*. In order to obtain the remaining sets of parameters, we use the intra-bunch distances (38). The mean intra-bunch distance allows us to extract the average time of transcriptional inactivity at the promoter which is equivalent to the time the promoter spends in the inactive state which does not lead to initiation. From Fig. 3A of reference (38) we extract the mean gap between RNAP bunches to be approximately 5 seconds. For the dead-end complex and ppGpp model, this implies that the average residence time of the promoter in 3^rd^ state is ~ 1/5=0.2/second. Hence for the three bursting models we take *k_OFF_* (dead-end complex model) = *k_OFF_* (ppGpp model) =*k_LOAD_^LOW^* (supercoiling mediated recruitment) ≈ 0.2/second. To obtain the fifth parameter of these models, we assume that the addition of ribosomal genes to the *E. coli* cell adjusts the overall transcription rate of the ribosomal genes by reducing the average transcription rate per gene, to keep the level of ribosomal RNA in the cell constant. In other words, for the models considered above the total initiation rate remains constant, or *nl = Constant*, where *n* is the number of ribosomal genes and *I* is the initiation rate on one of the genes. The initiation rate for a wild type pBR322 strain with *n* = 7 genes is *I* = 1 initiation/second(38). Using the formulas, we obtain for the rate of average initiation for each of the models of transcription initiation (see the SI), we find for the supercoiling mediated recruitment model *k_RELAX_* ≈ 0.055/second and for the *k_DEAD_* (dead-end complex) = *k_ON_* (ppGpp model) ≈ 0.047/second.

### Moments of intra-bunch distance distributions

To test the first and second class of models (along with the “free RNAP hypothesis”), we use the condition *nl* = Constant (i.e. the total number of ribosomal RNAs remain constant when the total gene number is increased) and change the rate of loading of RNAP molecules on to the promoter with the increasing number of genes, to keep the total initiation rate fixed. For each of these gene numbers we calculate the distribution of inter-polymerase distances along the gene. From this distribution, we construct the distributions of distances between RNAPs within a bunch and between bunches and compute their means and variances. In Fig. 4 we show the statistics of intra-bunch distances. For the third class of models, we change *k_ON_* as the amount of ppGpp molecules have been observed to increase with increasing gene numbers (73), and repeat the same exercise as before (also shown in Fig. 4 and Fig. S3 respectively)

## Acknowledgements

We wish to thank Rob Phillips, Hernan Garcia, Jeff Gelles and Timothy Harden for years of stimulating discussions and shared thoughts about transcriptional dynamics.

## References

1. 2007. Comparative Genomics: Volumes 1 and 2. Totowa (NJ): Humana Press.

2. Varki, A., and T.K. Altheide. 2005. Comparing the human and chimpanzee genomes: Searching for needles in a haystack. Genome Res. 15: 1746–1758.

3. Cheng, Y., Z. Ma, B.-H. Kim, W. Wu, P. Cayting, A.P. Boyle, V. Sundaram, X. Xing, N. Dogan, J. Li, G. Euskirchen, S. Lin, Y. Lin, A. Visel, T. Kawli, X. Yang, D. Patacsil, C.A. Keller, B. Giardine, mouse ENCODE Consortium, A. Kundaje, T. Wang, L.A. Pennacchio, Z. Weng, R.C. Hardison, and M.P. Snyder. 2014. Principles of regulatory information conservation between mouse and human. Nature. 515:371–375.

4. Stergachis, A.B., S. Neph, R. Sandstrom, E. Haugen, A.P. Reynolds, M. Zhang, R. Byron, T. Canfield, S. Stelhing-Sun, K. Lee, R.E. Thurman, S. Vong, D. Bates, F. Neri, M. Diegel, E. Giste, D. Dunn, J. Vierstra, R.S. Hansen, A.K. Johnson, P.J. Sabo, M.S. Wilken, T.A. Reh, P.M. Treuting, R. Kaul, M. Groudine, M.A. Bender, E. Borenstein, and J.A. Stamatoyannopoulos. 2014. Conservation of transacting circuitry during mammalian regulatory evolution. Nature. 515: 365–370.

5. Lin, S., Y. Lin, J.R. Nery, M.A. Urich, A. Breschi, C.A. Davis, A. Dobin, C. Zaleski, M.A. Beer, W.C. Chapman, T.R. Gingeras, J.R. Ecker, and M.P. Snyder. 2014. Comparison of the transcriptional landscapes between human and mouse tissues. Proc. Natl. Acad. Sci. U. S. A. 111: 17224–17229.

6. Loots, G.G. 2008. Genomic Identification of Regulatory Elements by Evolutionary Sequence Comparison and Functional Analysis. Adv. Genet. 61: 269–293.

7. Alberts, B., A. Johnson, J. Lewis, M. Raff, K. Roberts, and P. Walter. 2002. Molecular Biology of the Cell. 4th ed. Garland Science.

8. Bintu, L., N.E. Buchler, H.G. Garcia, U. Gerland, T. Hwa, J. Kondev, T. Kuhlman, and R. Phillips. 2005. Transcriptional regulation by the numbers: applications. Curr. Opin. Genet. Dev. 15:125–135.

9. Garcia, H.G., A. Sanchez, T. Kuhlman, J. Kondev, and R. Phillips. 2010. Transcription by the Numbers Redux: Experiments and Calculations that Surprise. Trends Cell Biol. 20: 723–733.

10. Garcia, H.G., and R. Phillips. 2011. Quantitative dissection of the simple repression input-output function. Proc. Natl. Acad. Sci. U. S. A. 108:12173–12178.

11. Rydenfelt, M., R.S. Cox, H. Garcia, and R. Phillips. 2014. Statistical mechanical model of coupled transcription from multiple promoters due to transcription factor titration. Phys. Rev. E. 89: 012702.

12. Zenklusen, D., D.R. Larson, and R.H. Singer. 2008. Single-RNA counting reveals alternative modes of gene expression in yeast. Nat. Struct. Mol. Biol. 15: 1263–1271.

13. Castelnuovo, M., S. Rahman, E. Guffanti, V. Infantino, F. Stutz, and D. Zenklusen. 2013. Bimodal expression of PHO84 is modulated by early termination of antisense transcription. Nat. Struct. Mol. Biol. 20: 851–858.

14. Raj, A., and A. van Oudenaarden. 2009. Single-Molecule Approaches to Stochastic Gene Expression. Annu. Rev. Biophys. 38: 255–270.

15. Jones, D.L., R.C. Brewster, and R. Phillips. 2014. Promoter architecture dictates cell-to-cell variability in gene expression. Science. 346:1533–1536.

16. Gandhi, S.J., D. Zenklusen, T. Lionnet, and R.H. Singer. 2011. Transcription of functionally related constitutive genes is not coordinated. Nat. Struct. Mol. Biol. 18: 27–34.

17. Raj, A., C.S. Peskin, D. Tranchina, D.Y. Vargas, and S. Tyagi. 2006. Stochastic mRNA synthesis in mammalian cells. PLoS Biol. 4: e309.

18. Padovan-Merhar, O., G.P. Nair, A.G. Biaesch, A. Mayer, S. Scarfone, S.W. Foley, A.R. Wu, L.S. Churchman, A. Singh, and A. Raj. 2015. Single Mammalian Cells Compensate for Differences in Cellular Volume and DNA Copy Number through Independent Global Transcriptional Mechanisms. Mol. Cell. 58: 339–352.

19. Golding, I., J. Paulsson, S.M. Zawilski, and E.C. Cox. 2005. Real-time kinetics of gene activity in individual bacteria. Cell. 123: 1025–1036.

20. Chubb, J.R., T. Trcek, S.M. Shenoy, and R.H. Singer. 2006. Transcriptional pulsing of a developmental gene. Curr. Biol. CB. 16:1018–1025.

21. Kepler, T.B., and T.C. Elston. 2001. Stochasticity in transcriptional regulation: origins, consequences, and mathematical representations. Biophys. J. 81: 3116–3136.

22. Sanchez, A., S. Choubey, and J. Kondev. 2013. Stochastic models of transcription: from single molecules to single cells. Methods San Diego Calif. 62: 13–25.

23. Bothma, J.P., H.G. Garcia, E. Esposito, G. Schlissel, T. Gregor, and M. Levine. 2014. Dynamic regulation of eve stripe 2 expression reveals transcriptional bursts in living Drosophila embryos. Proc. Natl. Acad. Sci. U. S. A. 111: 10598–10603.

24. Das, D., S. Dey, R.C. Brewster, and S. Choubey. 2017. Effect of transcription factor resource sharing on gene expression noise. PLoS Comput. Biol. 13: e1005491.

25. Mitarai, N., I.B. Dodd, M.T. Crooks, and K. Sneppen. 2008. The generation of promoter-mediated transcriptional noise in bacteria. PLoS Comput. Biol. 4: e1000109.

26. Elgart, V., T. Jia, A.T. Fenley, and R. Kulkarni. 2011. Connecting protein and mRNA burst distributions for stochastic models of gene expression. Phys. Biol. 8: 046001.

27. Singh, A., B. Razooky, C.D. Cox, M.L. Simpson, and L.S. Weinberger. 2010. Transcriptional Bursting from the HIV-1 Promoter Is a Significant Source of Stochastic Noise in HIV-1 Gene Expression. Biophys. J. 98: L32–L34.

28. Dar, R.D., B.S. Razooky, A. Singh, T.V. Trimeloni, J.M. McCollum, C.D. Cox, M.L. Simpson, and L.S. Weinberger. 2012. Transcriptional burst frequency and burst size are equally modulated across the human genome. Proc. Natl. Acad. Sci. 109: 17454–17459.

29. Kumar, N., A. Singh, and R.V. Kulkarni. 2015. Transcriptional Bursting in Gene Expression: Analytical Results for General Stochastic Models. PLoS Comput. Biol. 11: e1004292.

30. Munsky, B., G. Neuert, and A. van Oudenaarden. 2012. Using Gene Expression Noise to Understand Gene Regulation. Science. 336: 183–187.

31. Platini, T., T. Jia, and R.V. Kulkarni. 2011. Regulation by small RNAs via coupled degradation: Mean-field and variational approaches. Phys. Rev. E. 84: 021928.

32. Dong, G.Q., and D.R. McMillen. 2008. Effects of protein maturation on the noise in gene expression. Phys. Rev. E. 77: 021908.

33. Singh, A., and P. Bokes. 2012. Consequences of mRNA Transport on Stochastic Variability in Protein Levels. Biophys. J. 103: 1087–1096.

34. Melamud, E., and J. Moult. 2009. Stochastic noise in splicing machinery. Nucleic Acids Res. 37: 4873–4886.

35. Baker, C., T. Jia, and R.V. Kulkarni. 2012. Stochastic modeling of regulation of gene expression by multiple small RNAs. Phys. Rev. E. 85: 061915.

36. Choubey, S., J. Kondev, and A. Sanchez. 2015. Deciphering Transcriptional Dynamics In Vivo by Counting Nascent RNA Molecules. PLOS Comput. Biol. 11: e1004345.

37. Xu, H., S.O. Skinner, A.M. Sokac, and I. Golding. 2016. Stochastic Kinetics of Nascent RNA. Phys. Rev. Lett. 117.

38. Voulgaris, J., S. French, R.L. Gourse, C. Squires, and C.L. Squires. 1999. Increased rrn gene dosage causes intermittent transcription of rRNA in Escherichia coli. J. Bacteriol. 181: 4170–4175.

39. French, S.L., Y.N. Osheim, F. Cioci, M. Nomura, and A.L. Beyer. 2003. In exponentially growing Saccharomyces cerevisiae cells, rRNA synthesis is determined by the summed RNA polymerase I loading rate rather than by the number of active genes. Mol. Cell. Biol. 23: 1558–1568.

40. French, S.L., M.L. Sikes, R.D. Hontz, Y.N. Osheim, T.E. Lambert, A.E. Hage, M.M. Smith, D. Tollervey, J.S. Smith, and A.L. Beyer. 2011. Distinguishing the Roles of Topoisomerases I and II in Relief of Transcription-Induced Torsional Stress in Yeast rRNA Genes. Mol. Cell. Biol. 31: 482–494.

41. French, S.L., and O.L. Miller. 1989. Transcription mapping of the Escherichia coli chromosome by electron microscopy. J. Bacteriol. 171: 4207–4216.

42. Gotta, S.L., O.L. Miller, and S.L. French. 1991. rRNA transcription rate in Escherichia coli. J. Bacteriol. 173: 6647–6649.

43. Condon, C., S. French, C. Squires, and C.L. Squires. 1993. Depletion of functional ribosomal RNA operons in Escherichia coli causes increased expression of the remaining intact copies. EMBO J. 12: 4305–4315.

44. Bertrand, E., P. Chartrand, M. Schaefer, S.M. Shenoy, R.H. Singer, and R.M. Long. 1998. Localization of ASH1 mRNA Particles in Living Yeast. Mol. Cell. 2: 437–445.

45. Yunger, S., L. Rosenfeld, Y. Garini, and Y. Shav-Tal. 2010. Single-allele analysis of transcription kinetics in living mammalian cells. Nat. Methods. 7: 631–633.

46. Larson, D.R., D. Zenklusen, B. Wu, J.A. Chao, and R.H. Singer. 2011. Real-Time Observation of Transcription Initiation and Elongation on an Endogenous Yeast Gene. Science. 332: 475–478.

47. Garcia, H.G., M. Tikhonov, A. Lin, and T. Gregor. 2013. Quantitative Imaging of Transcription in Living Drosophila Embryos Links Polymerase Activity to Patterning. Curr. Biol. 23: 2140–2145.

48. Lucas, T., T. Ferraro, B. Roelens, J. De Las Heras Chanes, A.M. Walczak, M. Coppey, and N. Dostatni. 2013. Live Imaging of Bicoid-Dependent Transcription in Drosophila Embryos. Curr. Biol. 23: 2135–2139.

49. Sanchez, A., and I. Golding. 2013. Genetic Determinants and Cellular Constraints in Noisy Gene Expression. Science. 342: 1188–1193.

50. Sanchez, A., S. Choubey, and J. Kondev. 2013. Regulation of Noise in Gene Expression. Annu. Rev. Biophys. 42: 469–491.

51. Sánchez, Á., and J. Kondev. 2008. Transcriptional control of noise in gene expression. Proc. Natl. Acad. Sci. 105: 5081–5086.

52. Rosenfeld, L., E. Kepten, S. Yunger, Y. Shav-Tal, and Y. Garini. 2015. Single-site transcription rates through fitting of ensemble-averaged data from fluorescence recovery after photobleaching: A fat-tailed distribution. Phys. Rev. E. 92: 032715.

53. Klumpp, S., and T. Hwa. 2008. Stochasticity and traffic jams in the transcription of ribosomal RNA: Intriguing role of termination and antitermination. Proc. Natl. Acad. Sci. 105:18159–18164.

54. Dennis, P.P., M. Ehrenberg, D. Fange, and H. Bremer. 2009. Varying Rate of RNA Chain Elongation during rrn Transcription in Escherichia coli. J. Bacteriol. 191: 3740–3746.

55. Muthukrishnan, A.-B., M. Kandhavelu, J. Lloyd-Price, F. Kudasov, S. Chowdhury, O. Yli-Harja, and A.S. Ribeiro. 2012. Dynamics of transcription driven by the tetA promoter, one event at a time, in live Escherichia coli cells. Nucleic Acids Res. 40: 8472–8483.

56. Paulsson, J. 2004. Summing up the noise in gene networks. Nature. 427: 415–418.

57. Iyer-Biswas, S., F. Hayot, and C. Jayaprakash. 2009. Stochasticity of gene products from transcriptional pulsing. Phys. Rev. E Stat. Nonlin. Soft Matter Phys. 79: 031911.

58. Kepler, T.B., and T.C. Elston. 2001. Stochasticity in transcriptional regulation: origins, consequences, and mathematical representations. Biophys. J. 81: 3116–3136.

59. Peccoud, J., and B. Ycart. 1995. Markovian Modeling of Gene-Product Synthesis. Theor. Popul. Biol. 48:222–234.

60. Hornos, J.E.M., D. Schultz, G.C.P. Innocentini, J. Wang, A.M. Walczak, J.N. Onuchic, and P.G. Wolynes. 2005. Self-regulating gene: an exact solution. Phys. Rev. E Stat. Nonlin. Soft Matter Phys. 72:051907.

61. Gillespie, D.T. 1977. Exact stochastic simulation of coupled chemical reactions. J. Phys. Chem. 81: 2340–2361.

62. Dennis, P.P., M. Ehrenberg, and H. Bremer. 2004. Control of rRNA Synthesis in Escherichia coli: a Systems Biology Approach. Microbiol. Mol. Biol. Rev. 68: 639–668.

63. Baracchini, E., and H. Bremer. 1991. Control of rRNA synthesis in Escherichia coli at increased rrn gene dosage. Role of guanosine tetraphosphate and ribosome feedback. J. Biol. Chem. 266: 11753–11760.

64. Bremer, H., P. Dennis, and M. Ehrenberg. 2003. Free RNA polymerase and modeling global transcription in Escherichia coli. Biochimie. 85: 597–609.

65. Stepanova, E., J. Lee, M. Ozerova, E. Semenova, K. Datsenko, B.L. Wanner, K. Severinov, and S. Borukhov. 2007. Analysis of promoter targets for Escherichia coli transcription elongation factor GreA in vivo and in vitro. J. Bacteriol. 189: 8772–8785.

66. Susa, M., T. Kubori, and N. Shimamoto. 2006. A pathway branching in transcription initiation in Escherichia coli. Mol. Microbiol. 59: 1807–1817.

67. Kubori, T., and N. Shimamoto. 1996. A Branched Pathway in the Early Stage of Transcription byEscherichia coliRNA Polymerase. J. Mol. Biol. 256: 449–457.

68. Liu, L.F., and J.C. Wang. 1987. Supercoiling of the DNA template during transcription. Proc. Natl. Acad. Sci. U. S. A. 84: 7024–7027.

69. Lim, H.M., D.E.A. Lewis, H.J. Lee, M. Liu, and S. Adhya. 2003. Effect of varying the supercoiling of DNA on transcription and its regulation. Biochemistry (Mosc.). 42:10718–10725.

70. Opel, M.L., and G.W. Hatfield. 2001. DNA supercoiling-dependent transcriptional coupling between the divergently transcribed promoters of the ilvYC operon of Escherichia coli is proportional to promoter strengths and transcript lengths. Mol. Microbiol. 39: 191–198.

71. Maitra, A., I. Shulgina, and V.J. Hernandez. 2005. Conversion of active promoter-RNA polymerase complexes into inactive promoter bound complexes in E. coli by the transcription effector, ppGpp. Mol. Cell. 17:817–829.

72. Barker, M.M., T. Gaal, C.A. Josaitis, and R.L. Gourse. 2001. Mechanism of regulation of transcription initiation by ppGpp. I. Effects of ppGpp on transcription initiation in vivo and in vitro. J. Mol. Biol. 305: 673–688.

73. Gourse, R.L., Y. Takebe, R.A. Sharrock, and M. Nomura. 1985. Feedback regulation of rRNA and tRNA synthesis and accumulation of free ribosomes after conditional expression of rRNA genes. Proc. Natl. Acad. Sci. U. S. A. 82:1069–1073.

74. So, L., A. Ghosh, C. Zong, L.A. Sepúlveda, R. Segev, and I. Golding. 2011. General properties of transcriptional time series in Escherichia coli. Nat. Genet. 43: 554–560.

75. Taniguchi, Y., P.J. Choi, G.-W. Li, H. Chen, M. Babu, J. Hearn, A. Emili, and X.S. Xie. 2010. Quantifying E. coli proteome and transcriptome with single-molecule sensitivity in single cells. Science. 329: 533–538.

76. Jia, T., and R.V. Kulkarni. 2010. Post-transcriptional regulation of noise in protein distributions during gene expression. Phys. Rev. Lett. 105: 018101.

77. Huh, D., and J. Paulsson. 2011. Random partitioning of molecules at cell division. Proc. Natl. Acad. Sci. U. S. A. 108: 15004–15009.

78. Huh, D., and J. Paulsson. 2011. Non-genetic heterogeneity from random partitioning at cell division. Nat. Genet. 43: 95–100.

79. Schmidt, U., E. Basyuk, M.-C. Robert, M. Yoshida, J.-P. Villemin, D. Auboeuf, S. Aitken, and E. Bertrand. 2011. Real-time imaging of cotranscriptional splicing reveals a kinetic model that reduces noise: implications for alternative splicing regulation. J. Cell Biol. 193: 819–829.

80. Churchman, L.S., and J.S. Weissman. 2012. Native elongating transcript sequencing (NET-seq). Curr. Protoc. Mol. Biol. Ed. Frederick M Ausubel Al. Chapter 4: Unit 4.14.1–17.

81. Shawn C Little, M.T. 2013. Precise developmental gene expression arises from globally stochastic transcriptional activity. Cell. 154: 789–800.

82. Ring, B.Z., W.S. Yarnell, and J.W. Roberts. 1996. Function of E. coli RNA Polymerase σ Factor-σ70 in Promoter-Proximal Pausing. Cell. 86: 485–493.

